# Before the Black Box

**DOI:** 10.1101/2025.04.14.648839

**Authors:** Caitlyn L Perry, Charles Robin

## Abstract

Ecdysteroid hormones are responsible for regulating arthropod moulting, but unlike other taxa which used sterol-derived hormones (e.g. vertebrate sex-determining hormones), arthropods cannot synthesis sterols *de novo*. Ecdysteroid synthesis is thus dependent on funnelling diverse dietary sterols into the conserved reactions of ecdysteroidogenesis. While the later steps of ecdysteroid synthesis are dominated by cytochrome P450s, earlier stages of this pathway (i.e. those acting in or before the ‘Black Box’ in which 7-dehydrocholesterol is converted to 5,22,25-trideoxyecdysone) involve more diverse enzymes. While numerous comparative analyses of arthropod cytochrome P450s have been conducted, little comparable work has been done on the pre-Black Box FAD-dependent oxidoreductase DHCR24 and the Rieske domain oxygenase Neverland and the Black Box short-chain dehydrogenase/reductase Shroud. However, there is some evidence for evolutionary shifts among these enzymes across arthropods, e.g. the absence of DHCR24 from *Drosophila melanogaster*. We have sought to systematically identify such changes in copy number in DHCR24, Neverland and Shroud across arthropods, as well as to identify correlations between these changes and dietary factors with reference to the literature.

Among other results, we find that Neverland is absent from all beetles (providing support for mycophagy as an important element of the beetle ancestral diet) and that losses of DHCR24 correlate with production of alkylated ecdysteroids.

## 1. Introduction

The ecdysteroids, best known as regulators of arthropod moulting, are distinctive among animal steroid hormones in their extensive hydroxylation. This structural property has important functional implications, such as increased hydrophilicity (with consequent reduction of cell membrane permeability; Okamoto *et al*., 2018), and it is therefore unsurprising that much literature describing the synthesis of ecdysteroids has focused on the cytochrome P450s responsible for these hydroxylation reactions (e.g. Rewitz *et al*., 2006; Iga and Kataoka, 2010). However, hydroxylation is not the only transformation needed to convert dietary sterols to ecdysteroids, and an understanding of the upstream reactions – including side-chain dealkylation, oxidation of bonds within the sterol rings and reduction of hydroxyl groups – is particularly important from an evolutionary perspective. While the cytochrome P450s responsible for ecdysteroid hydroxylation are, with a single exception (Nelson, 2018), restricted to arthropods, other ecdysteroidogenic enzymes (namely DHCR24, Neverland and Shroud) have more ancient roles in steroid metabolism.

Research into ecdysteroid synthesis has tended to assume cholesterol as the starting point, but only exclusively carnivorous arthropods consume no other sterol in relevant quantities. Campesterol, stigmasterol, beta–sitosterol and ergosterol are among the sterols most frequently consumed by arthropods feeding on plant or fungal tissue, and these molecules share a distinctive feature – the presence of an alkyl side-chain at C24. When these molecules are used as substrates for ecdysteroid synthesis, the side-chains may be retained or removed; both possibilities are realised across disparate insect species. Although the reactions involved in the transformation of plant and fungal sterols to cholesterol are well established at least for some species (e.g. Svoboda *et al*., 1975), little progress has been made in identifying the enzymes which catalyse them. There is evidence that a cytochrome P450 is responsible for fucosterol-(24,28)-epoxide cleavage in *Spodoptera littoralis* (Clarke *et al*., 1985), and cytochrome P450s have also been associated with earlier stages of sterol dealkylation (Gilbert *et al*., 2002), but as most insects have a sizeable complement of cytochrome P450s (e.g. 120 in *S. littoralis* – Wu *et al*., 2022), identification of side-chain cleavage enzymes among them is not trivial. The final step of side-chain dealkylation, removal of the C24(25) double bond, is by contrast well understood. DHCR24 has been shown to catalyse this reaction in *Bombyx mori* (Cuifo *et al*., 2011), and its orthologs play the same role in other animals, though in a different context. Desmosterol-24-reduction is ancestrally a part of cholesterol synthesis; why it was retained in arthropods despite the otherwise complete loss of the biosynthetic pathway responsible for converting squalene to cholesterol (Clark and Bloch, 1959) is unclear, though the presence of sterols with C24 side-chains in the ancestral arthropod diet would provide an explanation.

Like DHCR24, Neverland has an ancestral role in cholesterol metabolism – specifically, it converts cholesterol to 7-dehydrocholesterol, the precursor of vitamin D (Prabhu *et al*., 2016), thus reversing the final step of cholesterol synthesis. In some lineages, notably mammals, this role has proven dispensable, with cholesterol-induced DHCR7 degradation instead serving to balance the two molecules. In the arthropod context, Neverland typically catalyses the conversion of cholesterol to 7-dehydrocholesterol, but deviation from this canonical role has been observed in *Drosophila pachea*, a species long known to depend either on extracts from its host senita cactus or on supplementation of Δ7 sterols such as lathosterol for growth (Heed and Kircher, 1965; Goodnight and Kircher, 1971). This requirement stems from a change in the substrate preference and catalytic activity of Neverland in the *D. pachea* lineage. The ancestral function of Neverland is obviously irrelevant to the metabolism of Δ7 sterols; however, some Δ7 sterols (including lathosterol and schottenol, both found in senita cactus) lack the C5(6) double bond present in cholesterol. In response to this unusual dietary sterol intake, *D. pachea* Neverland has specialised for sterol-5-reductase activity to such an extent as to have lost sterol-C7-reductase activity entirely, with four amino acid substitutions playing major roles in this shift (Lang *et al*., 2012). While the example of *D. pachea* provides evidence for some lability in the role of Neverland, it is not yet clear just how exceptional this case is.

The role of Shroud in the synthesis of ecdysteroids was established by Niwa *et al*. (2010); these authors assigned the enzyme to the Black Box on the grounds that RNA interference against *D. melanogaster shroud* could be rescued by feeding with 2,22,25-trideoxyecdysone but not 7-dehydrocholesterol. However, this publication did not speculate as to the precise catalytic role of Shroud, a topic which has in fact received very little discussion in the literature in general. As Shroud is a short-chain dehydrogenase/reductase, it seems very likely that it catalyses one of the two reductions which occur within the Black Box (i.e. 3-hydroxyl-reduction or 14-hydroxyl-reduction); the former reaction has a parallel in the synthesis of dafachronic acids from sterols in *Caenorhabditis elegans* (and presumably other nematodes), in which 3-hydroxyl reduction is catalysed by the short-chain dehydrogenase/reductase DHS-16 (Wollam *et al*., 2012).

In order to deepen our understanding of ecdysteroid evolution and its connection to factors such as diet, we have analysed the distribution, copy number and (where relevant) sequence evolution of these enzymes across 1432 publicly available arthropod genome assemblies. We have also performed some analyses touching on other enzyme families implicated in the conversion of phytoecdysteroids into sterols, namely cytochrome P450s and GMC oxidoreductases, though our ignorance of the specific enzymes catalysing these reactions in any species imposes limitations here.

## 2. Methods

The amino acid sequences of *Drosophila melanogaster* Shroud and Neverland and of *Bombyx mori* DHCR24 (as *D. melanogaster* belongs to a lineage which has lost DHCR24) were used as queries in TBLASTN orthology searches with default NCBI server settings across 1432 genome assemblies (containing one member of all arthropod genera represented in GenBank as of June 2023). Where multiple representatives of a genus existed at that time, the assembly with the highest N50 value was examined; if that assembly indicated duplication or loss of the query gene, other representatives of the genus were also examined. Multiple hits in a genome showing comparable similarity to the query sequence were taken to reflect gene duplicates, while the absence of a reciprocal best hit was treated as a gene loss. Transcriptomes and raw sequencing data were used sporadically to clarify the timing of gene duplications and losses where genome assembles were not available. To reduce the risk of contaminating DNA distorting our results, hits from transcriptomes or raw genomic sequences were subject to reciprocal BLAST search and retained only if the best hits came from closely related taxa.

Amino acid sequences used for phylogenetic inference (e.g. in determining the number of duplications which have occurred in a particular lineage) were obtained from transcriptome data, automated gene prediction (with manual checking) or manual annotation. Sequences were aligned using MAFFT with L–INS–i settings (Katoh and Toh, 2008). IQ–TREE version 1.6.1 (Nguyen *et al*., 2015) was used to model sequence evolution, with 1000 ultrafast bootstrap replicates and 1000 SH-like approximate likelihood ratio replicates produced (Hoang *et al*., 2018). Amino acid substitution models were chosen by ModelFinder (Kalyaanamoorthy *et al*., 2017) with default settings based on BIC value. Phylogenetic trees were rendered with TreeViewer (www.treeviewer.org), with nodes polytomised where SH-aLRT support was below 80 or ultrafast bootstrap support was below 95.

We found that one–to–one orthologs of Shroud could not be reliably detected outside Hexapoda by reciprocal BLAST search. In order to test for Shroud orthologs in other arthropods, the alignment and phylogenetic inference methods described above were used to produce phylogenies of short– chain reductases for representatives of crustaceans and chelicerates (no available myriapod transcriptome contained enough full–length hits to justify confidence that these represented even a mostly complete inventory of short–chain reductases). Twenty *D. melanogaster* short–chain reductases (including CG8888, described by Niwa *et al*. (2010) as closely related to Shroud) were aligned with all detected single–domain short–chain reductases from the species *Hyalella azteca*, *Triops cancriformis*, *Tigriopus japonicus*, *Amphibalanus amphitrite*, *Argulus siamensis* and *Parasteatoda tepidariorum*. Crustacean and chelicerate short-chain reductases were identified on the basis of a hit (at the default E < 0.05 level) in a TBLASTN search with *D. melanogaster* Shroud as the query and the presence of only a short–chain reductase domain in a Conserved Domain search (Marchler–Bauer *et al*., 2017) with default NCBI server settings. Alignment and tree inference were then performed as described above. One-to-one orthologs of *D. melanogaster* Shroud identified through this procedure were used as query terms in future searches in the relevant group.

To examine changes in selective pressure (e.g. to determine which of multiple orthologs of a *D. melanogaster* ecdysteroid synthesis enzyme is most likely to have retained ancestral function), amino acid alignments were converted into DNA alignments using RevTrans v.2 with default settings (Wernersson and Pedersen, 2003). These alignments were used to compare of synonymous and non–synonymous substitution ratios using Contrast-FEL, a component of the HyPhy package (Kosakovsky Pond and Muse, 2005; Kosakovsky Pond *et al*., 2021; all programs used with default settings). A second HyPhy package, RELAX (Wertheim *et al*., 2015), was used with to analyse relaxation of selective constraint in aedine mosquito DHCR24, while evolution of the *Helicoverpa armigera* and *H. zea* CYP4G15 relatives (with *D. melanogaster* CYP4G15 used as an outgroup) was examined with aBSREL (Smith *et al*., 2015).

To identify Rieske short-chain oxygenases in *Tribolium castaneum*, we used the Pfam PF00355 profile to query the *T. castaneum* protein set (GCF_031307605.1) with the hmmsearch command in HMMER 3.1 (hmmer.org, default settings except that heuristic filters were turned off).

## 3. Results and Discussion

### 3.1. DHCR24 Copy Number Variation

#### 3.1.1. DHCR24 Losses

At least fifteen losses of DHCR24 are apparent among examined arthropod genomes, and a strong correlation between retention of DHCR24 and capacity for dealkylation of phyto- and mycosterols is apparent (Table 1). This relationship is consistent with expectations based on DHCR24’s documented role in sterol dealkylation in *Bombyx mori* and in vertebrates. While we have data on sterol side-chain dealkylation for only a small number of species, several species which have been demonstrated to retain sterol side-chains (e.g. *Apis mellifera*, *Oncopeltus fasciatus*) lack DHCR24, while several which are known to remove side-chains possess it (e.g. *Blattella germanica*, *Manduca sexta*). *Solenopsis invicta* is a possible counterexample, but the evidence for dealkylation in this species is limited to the observation that cholesterol makes up around half the total sterol present in individuals of this species at several developmental stages despite the lack of an obvious source of dietary cholesterol (Ba *et al*., 1995). However, given the capacity of many insects to selectively uptake particular sterols (e.g. Carvalho *et al*., 2012), low dietary cholesterol alone is insufficient grounds for assuming an organism can dealkylate phytosterols. It is of course possible that groups lacking DHCR24 might develop a novel mechanism for removing the C23(24) double bond, or even a means of C24-dealkyation which does not generate this double bond. In general, however, absence of DHCR24 from a good-quality genome assembly is a strong predictor of inability to dealkylate sterols.

**Table 1.** Groups which have lost DHCR24 and the ecdysteroids produced by their representatives (where known).

| Taxon | Ecdysteroids produced |
| --- | --- |
| Adephaga (Coleoptera) | Unknown |
| Cantharidae+Lampyridae (Coleoptera) | Unknown |
| Acariformes | Ponasterone A (Grbić *et al.*, 2011) – dealkylated, but lacking 25-hydroxyl group |
| Pseudoscorpiones | Unknown |
| Tetrapulmonata | Unknown |
| Chironomoidea (Diptera) | Unknown |
| Brachycera (Diptera) | Makisterone A in *Drosophila melanogaster* (Redfern, 1984) and *Musca domestica* (Feldlaufer and Svoboda, 1991) |
| Coccoidea (Hemiptera) | Unknown |
| Heteroptera (Hemiptera) | Makisterone A in *Megalotomus quinquespinosus* (Feldlaufer *et al.*, 1986), *Dysdercus cingulatus*, *Nezara viridula*, *Podisus maculiventris* (Aldrich *et al.*, 1982; Kelly *et al.*, 1984), makisterone A and C in *Dysdercus fasciatus* (Feldlaufer *et al.*, 1991) |
| Aculeata (Hymenoptera) | Makisterone A in *Apis mellifera* (Feldlaufer *et al.*, 1985), 24-epimakisterone A in *Acromyrmex octospinosus* (Maurer *et al.*, 1993) |
| Chrysidoidea (Hymenoptera) | Unknown |
| Ichneumonoidea (Hymenoptera) | Unknown |
| Platygastroidea (Hymenoptera) | Unknown |
| Odonata | Unknown |
| Neuropterida (Raphidioptera uncertain) | Unknown |

We can easily imagine evolutionary scenarios in which inability to dealkylate phyto- and mycosterols would impose no selective disadvantage; wherever cholesterol (or another sterol lacking a C24 side chain, e.g. lathosterol) is abundant in the diet, maintenance of sterol dealkylation machinery is unnecessary. Some observed losses of DHCR24 correlate pleasingly with established dietary changes; for example, the suborder Adephaga, unusual among beetles in lacking DHCR24, has been established as ancestrally predacious (Beutel *et al*., 2020). While multiple separate DHCR24 losses have evidently occurred among Hymenoptera, all affected lineages sit within an ancestrally parasitoid clade (Peters *et al*., 2017). Heteroptera, one of two hemipteran lineages which clearly lacks DHCR24, is generally considered ancestrally predacious, despite a degree of uncertainty about the relationships among basal heteropterans (Li *et al*., 2017). However, some equally strongly supported DHCR24 losses are more difficult to explain through dietary factors, as with the other hemipteran group lacking DHCR24, Coccoidea (scale insects). Scale insects are exclusively phytophagous; more specifically, they exclusively consume plant phloem, which is very low in sterols. Yeast-like endosymbionts appear to provide sterols to at least some scale insects (Eya *et al*., 1989); ergosterol is the major product of these endosymbionts under aerobic conditions, though ergosta-5,7,24(28)-trienol has been retrieved as their main sterol under anaerobic conditions, i.e. *in vivo* (Wetzel *et al*., 1992). ERG5 is expressed at very low levels in yeast-like endosymbionts of planthoppers, consistent with an abundance of ergosta-5,7,24(28)-trienol (Noda and Koizumi, 2003). However, whether the predominant sterol of yeast-like endosymbionts is ergosterol or ergosta-5,7,24(28)-trienol, DHCR24 would be expected to be required for side-chain cleavage.

It is possible that DHCR24 was lost from Coccoidea as part of its transition to a low-sterol diet, prior to the development of endosymbionts capable of providing sterols to their hosts, but we should not be rigidly wedded to dietary explanations of DHCR24 losses in all cases. For example, the diets of anopheline and culicine mosquitoes are very similar, leaving little room for any history of substantial differences in sterol intake to explain loss of DHCR24 in the former subfamily and retention in the latter. It may be that juvenile and adult female consumption of cholesterol in *Anopheles* species is sufficient to allow synthesis of 20-hydroxyecdysone at required levels without side-chain dealkylation; if so, *Aedes* DHCR24 should also be dispensable and thus under relaxed selective constraint. However, a comparison of five culicine mosquito DHCR24 sequences and all other available nematoceran DHCR24 sequences using RELAX in fact suggests higher selective constraint on mosquito DHCR24 (p=0.0015). Sterol dealkylation therefore appears functionally significant in modern culicine mosquitoes, although what that function might be remains unclear.

The possibility that synthesis of alkylated ecdysteroids (e.g. makisterone A rather than 20-hydroxyecdysone) has been selectively neutral or even advantageous in a given lineage should be considered in trying to account for puzzling DHCR24 distributions. Makisterone A appears to be a weaker activator of the EcR-Ultraspiracle receptor than 20-hydroxyecdysone in most species (though exceptions in phytophagous lineages lacking DHCR24 are known, e.g. Tohidi-Esfahani *et al*., 2011), but we can imagine circumstances where reduced hormonal activity might be beneficial, for example if a dietary shift has resulted in increased sterol intake. Another possible explanation for DHCR24 loss is the phenomenon of selective cholesterol uptake mentioned above. While most plants and many fungi contain only trace amounts of cholesterol, preferential use of cholesterol over more abundant alkylated sterols could allow arthropods lacking DHCR24 to synthesise 20-hydroxyecdysone. Cholesterol storage in holometabolous insects may allow synthesis of 20-hydroxyecdysone even where cholesterol is absent from the adult diet; several studies (Pondeville *et al*., 2008; Peng *et al*., 2022) have observed synthesis of 20-hydroxyecdysone in *Anopheles* males with no dietary sterol intake. In laboratory settings, *Anopheles* larvae are generally fed with commercial fish food, which includes animal products (and therefore some cholesterol), explaining synthesis of 20-hydroxyecdysone rather than makisterone A etc. The sterol intake of wild *Anopheles* larvae may of course be quite different from their typical laboratory diet; determining the composition of the ecdysteroid titre in wild adult mosquitoes would be necessary to fully clarify the functional significance of DHCR24 loss in the *Anopheles* genus.

Finally, certain patterns in the distribution of DHCR24 remain difficult to interpret in evolutionary terms due to phylogenetic uncertainty. This is most obviously the case among Chelicerata, as the monophyly of terrestrial chelicerates appears increasingly implausible (Ballesteros *et al*., 2022). Table 1 follows the phylogeny of Ballesteros *et al*. in placing gene losses; the lifestyle and diet of the organisms in which they occurred are (at least currently) unknowable.

#### 3.1.2. DHCR24 Duplications

The initial description of DHCR24 in arthropods (Ciufo *et al*., 2011) noted the presence in many (though not all) species which had conserved DHCR24 of a paralog referred to as ‘DHCR24-like’. This enzyme is clearly distinct from DHCR24 at the sequence level; Ciufo *et al*. report amino acid identity levels between 34 and 42% (their gene structures are also divergent, with two introns in most DHCR24-like genes vs. nine in DHCR24). Phylogenetic inference (Figure 1) suggests that the split of DHCR24 and DHCR24-like occurred in a common ancestor of most or all crustaceans and insects; both *Daphnia magna* (a water flea) and *Capitulum mitella* (a barnacle) possess two genes related to DHCR24, one of which clearly groups with *Bombyx mori* DHCR24-like. While their paralogs do not cluster unambiguously with *B. mori* DHCR24, this likely reflects the relatively short branch lengths associated with these genes; arthropod clades which possess a single DHCR24 ortholog, e.g. myriapods and pycnogonids, likewise do not cluster clearly with either DHCR24 or DHCR24-like. Given the age of the split between DHCR24 and DHCR24-like, it is unlikely that the ancestral role of DHCR24-like lies in phytosterol dealkylation, as Ciufo *et al*. hypothesised, though this certainly does not exclude the possibility that it has acquired such a role in modern arthropod species. Transcriptome data could assist in clarifying this point, but the absence of DHCR24-like from many of the groups with the richest transcriptomic resources is currently an obstacle to such analyses.

**Figure 1.**
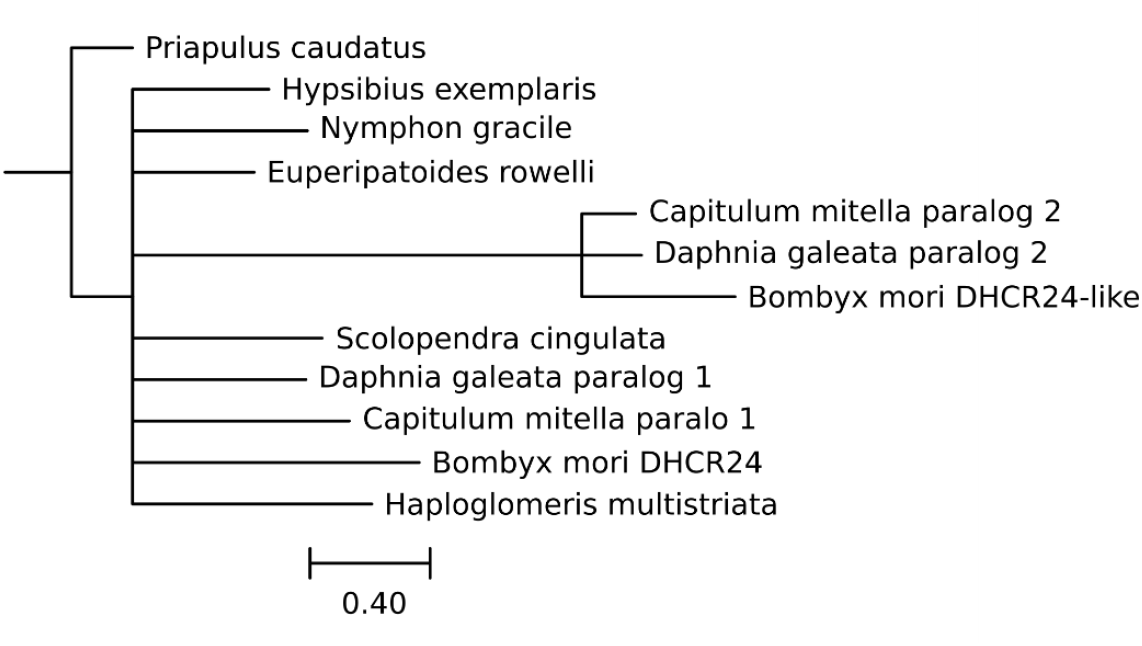
Maximum likelihood tree of DHCR24 and DHCR24-like orthologs. Species represented are the silkworm *Bombyx mori*, the water flea *Daphnia galeata*, the barnacle *Capitulum mitella*, the centipede *Scolopendra cingulata*, the millipede *Haploglomeris multistriata*, the pycnogonid *Nymphone gracile*, the velvet worm *Euperipatoides rowelli*, the tardigrade *Hypsibius exemplaris* and the penis worm *Priapulus caudatus* (outgroup). An *H. exemplaris* RNA sequence (GFGW01014089.1) with high identity to *B. mori* DHCR24 was excluded due to failure to map to the *H. exemplaris* genome assembly.

Duplications of DHCR24 itself are sporadic, but seem to occur more frequently in specific groups, namely the springtails (Collembola), Lepidoptera, Orthoptera, Aphidoidea and the fly superfamily Sciaroidea (although this interpretation may be influenced by uneven representation of arthropod taxa in public genome databases). In some cases, a single ancestral duplication is reflected in numerous species, as with Aphidoidea, while other groups show more complex patterns. An example comes from the moth subfamily Ennominae (Figure 2), where secondary losses (e.g. the retention of two DHCR24 paralogs in *Biston betularia* but not *B. stratarius*) and pseudogenisation (e.g. a partial sequence in *Alcis repandata* not represented in Figure 2) complicate the distribution. At least four duplications appear to have occurred in Papilionoidea – three are found only in a single genus (*Leptidea, Delias*, and *Calephelis*), while one is shared across the tribe Argynnini (Supplementary Figure 1). The distribution of DHCR24 homologs in Orthoptera is suggestive of an early duplication with subsequent loss in Caelifera, with *Vandiemenella viatica*’s two copies resulting from a secondary duplication (Supplementary Figure 2). Collembola shows by far the most striking DHCR24 radiations (most notably in *Mesaphorura yosii*), but most of these are confined to a single species (Figure 3), at least among those for which genome assemblies were available.

**Figure 2.**
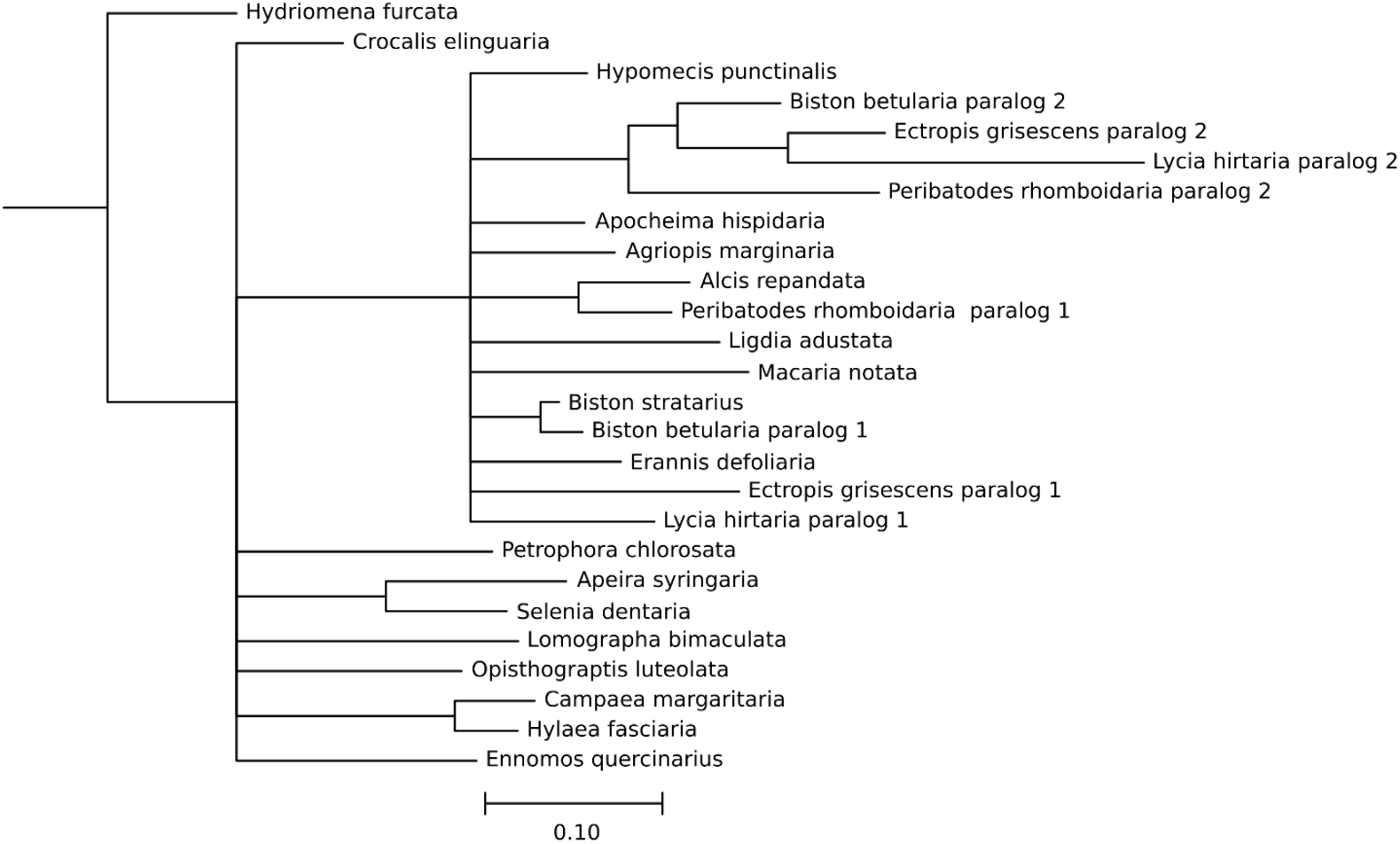
Maximum likelihood tree of DHCR24 and DHCR24-like orthologs from the moth subfamily Ennominae. All members of Ennominae in our dataset (Supplementary Table 1) from which full-length amino acid sequences could be determined are included. *Hydriomena furcata* (Larentiinae) DHCR24 was chosen as an outgroup on the basis that Larentiinae is well-supported as monophyletic relative to the geometrid clade containing Ennominae (Murillo-Ramos *et al*., 2019).

**Figure 3.**
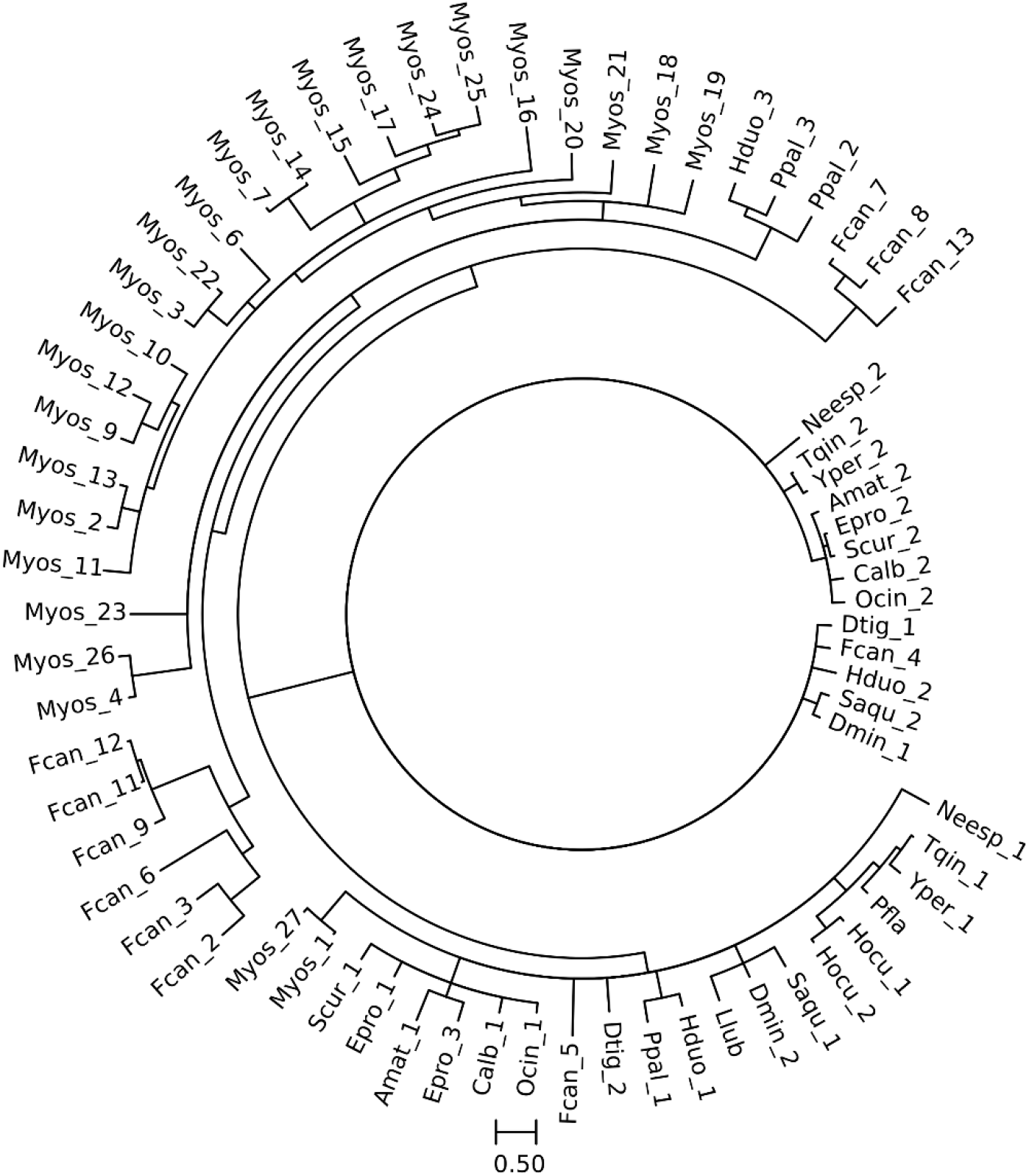
Maximum likelihood tree of DHCR24 orthologs from Collembola. Species in which the sequence of one or more DHCR24 orthologs could not be reconstructed fully due to sequencing gaps or apparent errors were removed, both for ease of interpretation and to avoid the implication of paralog losses which have probably not occurred; otherwise all collembolan species in our dataset (Supplementary File 1) are included.

Duplication of DHCR24 could potentially be explained by specialisation for activity on sterols with differing side chain structures, e.g. a single (campesterol, sitosterol) vs. double (ergosterol, stigmasterol, brassicasterol) bond at C22(23). However, evidence that the C22(23) double bond is removed before the C24(25) double bond at least in *Manduca sexta* (Svoboda *et al*., 1969) implies that this explanation is unlikely to be universally applicable. Of course, it is possible that one of the DHCR24 paralogs in fact acts as a sterol-22-reductase. Such a shift in function would be expected to involve positive selection at some sites; while a Contrast-FEL analysis did not reveal any sites under differential selection between the two clades in species with both paralogs, the small number of species under analysis likely limits our capacity to detect site-specific rate changes. Ultimately, experimental characterisation of DHCR24 paralogs will be necessary to determine whether these are typically functionally differentiated.

#### 3.1.3. Sterol Dealkylation – DHCR24 and Colleagues

Better knowledge of the enzymes acting upstream of DHCR24 would help clarify its evolutionary history, and one potential means of winnowing our candidate list is apparent – restricting the search to enzymes induced by consumption of sterols containing C24 alkyl groups. This possibility was one motivation for Jing *et al*.’s (2012) examination of the effects of stigmasterol vs. cholesterol feeding on gene expression in *Helicoverpa zea*. While DHCR24 was not explicitly discussed by the authors (not having been identified in insects at the time the research was conducted), examining their supplementary data reveals that it is not among the genes showing a greater than twofold difference in expression between these two feeding conditions. This result shows the limitations of expression-based analysis, but does not exclude the possibility of other enzymes required for sterol side-chain dealkylation being upregulated by their substrates. Jing *et al*. propose several candidate reductases, two of which belong to the GMC oxidoreductase cluster (Iida *et al*., 2007). One member of the equivalent cluster in *D. melanogaster* acts as an ecdysone-3-oxidase (Takeuchi *et al*., 2005; Iida *et al*., 2007); while ecdysone-3-oxidase reduces a carbon-oxygen rather than a carbon-carbon bond, and acts on a sterol ring rather than the side-chain, this similarity in substrates justifies further examination of the *H. zea* GMC cluster. The other proposed candidates lack equally clear *D. melanogaster* orthologs, complicating the identification of candidate substrates on the basis of homology.

Jing *et al*. also discuss several cytochrome P450s which might act as side-chain epoxidases; among these is an ortholog of *H. armigera* CYP340H1. This enzyme has not received detailed study, and although it is known to be overexpressed in a cypermethrin-resistant lineage, this association does not appear to be causal (Rasool *et al*., 2015). However, other enzymes in the CYP340 subfamily act as epoxidases in the context of lepidopteran pheromone synthesis (Rong *et al*., 2019). While this combination of expression data and functional analysis makes CYP340H1 a plausible sterol epoxidase in *H. armigera*, the CYP340 group is limited to Lepidoptera (Fujii *et al*., 2020), suggesting that steroid side-chain dealkylation may have been elaborated independently in the several arthropod lineages where it occurs. The other cytochrome P450s discussed by Jing *et al*. in the context of steroid side-chain epoxidation all belong to the CYP6 family, members of which are prominent in xenobiotic and insecticide detoxification, though the expression patterns of some CYP6 enzymes (e.g. Chung *et al*., 2009) suggest endogenous substrates. CYP6 enzymes can catalyse diverse reactions (see e.g. Hoi *et al*., 2014), but no epoxidase function for any enzyme of this subfamily has yet been established.

One additional cytochrome P450 should be discussed in the context of *H. zea* sterol dealkylation – CYP4G8, a CYP4G15-like enzyme upregulated not only in response to stigmasterol feeding (38-fold), but also to dietary cholestanol (45-fold) and cholestan-3-one (72-fold), raising the possibility that it may be downregulated by dietary cholesterol. The use of homology as a guide to the likely function of this enzyme is complicated by the intricacies of the CYP4G group’s evolution. CYP4G1 enzymes have been identified as necessary for the decarbonylation of fatty aldehydes in cuticular hydrocarbon synthesis (Qiu *et al*., 2012), as have some CYP4G15 enzymes (e.g. Chen *et al*., 2016). As CYP4G1 and CYP4G15 are sister clades (Feyereisen, 2020), catalysis of fatty aldehyde decarbonylation is likely to have been the original function of the common ancestor of the CYP4G1 and CYP4G1. The previously-observed relationship between CYP4G8 expression and pyrethroid resistance (Pittendrigh *et al*., 1997) thus likely reflects reduced insecticide uptake due to changes in cuticular chemistry (Xin *et al*., 2022). While the expression of *D. melanogaster Cyp4g15* in the embryonic, larval and adult central nervous system (Maϊbèche-Coisne *et al*., 2000; Chung *et al*., 2009) indicates that not all CYP4G1-4G15 enzymes act on cuticular components, there is no clear evidence for novel functional roles among the *Helicoverpa* ‘CYP4G15-like’ enzymes. Given the substantial structural differences between fatty aldehydes and steroids and evidence that the stigmasterol-induced ‘CYP4G15-like’ enzyme is not evolving especially rapidly (uncorrected *p*-value for this branch = 1 in aBSREL analysis), a role for this enzyme in phytosterol dealkylation seems highly unlikely, though the effect of dietary sterols on CYP4G8 expression remain to be explained.

We therefore cannot, at present, identify the enzymes acting prior to DHCR24 in the removal of the side-chain from plant sterols in *H. zea*. Even if we were able to do so, how much would this tell us about these reactions in other arthropods? It appears highly implausible that all the reactions of phytosterol dealkylation are conversed across arthropods, as any cytochrome P450 retained in most or all arthropods which dealkylate phytosterols, but generally lost by those which do not, should have been identified in large-scale analyses of insect cytochrome P450s such as Dermauw *et al*. (2020). Equally, one-to-one orthology in the GMC oxidoreductase cluster is imperfect even within insect orders (Iida *et al*., 2007), but there is no obvious sign of a GMC subfamily restricted to arthropods which dealkylate phytosterols. The suggestion that the enzymes catalysing the first three steps of sterol side-chain dealkylation have been subject to turnover where DHCR24 has been conserved may initially seem unconvincing, but it should be remembered that DHCR24 catalyses the only step of dealkylation in which the substrate is identical regardless of the sterol consumed (e.g. Svoboda and Weirich, 1995). Changes in the prevalence of, for instance, C28 vs. C29 sterols in the diet could serve as a selective pressure favouring the turnover of enzymes with newly unsuitable substrate preferences, but would not affect DHCR24.

While the instability of DHCR24 may surprise readers, the long tradition of treating ecdysteroid synthesis as beginning with cholesterol might encourage the perception of this enzyme as lying outside the ecdysteroid synthesis pathway proper, with a degree of phylogenetic unruliness thus to be expected. However, an examination of the better-credentialled ecdysteroidogenic enzymes, to which we now turn, reveals entirely comparable – and in many cases less easily explicable – instability in both copy number and function.

### 3.2. Neverland: Losses and Blooms

#### 3.2.1. Neverland Losses

The taxonomically broadest loss of Neverland extends across all beetles and possibly their sister group, the twisted-wing parasites (Strepsiptera). Strepsiptera was represented in our dataset by a single low-coverage genome assembly and a small number of transcriptomes; while the only Neverland hits from these sources can be attributed to host contamination on the basis of sequence similarity, the evidence base is still too thin to be entirely comfortable drawing firm conclusions. In any case, the loss cannot extend further than Strepsiptera, as members of Neuropterida, which is sister to the combined Coleoptera-Strepsiptera group, possess a Neverland ortholog. A possible driver of this gene loss can be identified by examining a smaller-scale Neverland loss within Sciaroidea, a group mentioned in section 3.1.2. in connection with DHCR24 duplication. Sciaroids are largely fungus feeders, and while ergosterol is by no means a singular ‘fungal sterol’ (Weete *et al*., 2010), it is nonetheless an extremely widespread sterol among fungi. As ergosterol contains both C5(6) and C7(8) double bonds, the sterol-C7-reductase activity of Neverland would be dispensable for an arthropod with high ergosterol intake.

While the great dietary diversity of modern beetles indicates that sterol-C7-reduction must occur in many beetle species, a dietary explanation for Neverland loss may apply to earlier stages in beetle evolution. An association between the earliest beetles and dead wood is generally accepted (e.g. Beutel *et al*., 2024), and several authors have suggested that the diet associated with this habitat was mycophagy rather than xylophagy (Crowson, 1981; Ponomarenko, 2003). As ergosterol synthesis occurs in both ascidiomycotes and basidiomycetes, which diverged 600-650 million years ago (Tedersoo *et al*., 2018), ergosterol would almost certainly have been the dominant sterol in the diet of putative mycophagous early beetles, given that the beetle radiation has been placed between 300 (Zhang *et al*., 2018) and 330 (Toussaint *et al*., 2016) million years ago. Whether the last common ancestor of Coleoptera and Strepsiptera was also fungus-feeding is of course a separate question. As the relationship with Coleoptera and Strepsiptera was only relatively recently established firmly (Niehuis *et al*., 2012), it is unsurprising that little research into their shared ancestors has been conducted. The loss of adult feeding in all strepsipterans, even the early-diverging Mengellidae, in which females are free-living (Tröger *et al*., 2019), makes reconstruction of their ancestral diet extremely challenging.

It should be emphasised at this point that ergosterol consumption is unlikely to explain all instances of Neverland loss. Potential counter-examples are provided by Neverland loss in the mite order Mesostigmata and the crustacean subclass Anostraca. Neither the ancestrally predacious lifestyle of the former group (Castilho *et al*., 2015) nor the filter-feeding of the latter (Reeve, 1963) can easily be reconciled with high ergosterol intake, though it is possible that some other Δ5,Δ7-sterol was abundant in the ancestral diet of one or both groups. Indeed, it is difficult to imagine specific factors which might drive loss of Neverland apart from diet, though this is not sufficient reason to rule out their existence.

Whatever evolutionary scenario led to the loss of Neverland from beetles (and perhaps strepsipterans), there is evidence that at least some modern beetles which do not consume ergosterol or other Δ5,Δ7-sterols nonetheless produce ecdysteroids with both double bonds (e.g. Delbecque and Sláma, 1980). It therefore seems likely that at least one sterol-C7-reductase has arisen within Coleoptera, although it is possible that symbiotic organisms could fill this role, especially given evidence for sterol provisioning by yeast-like symbiotes in anobiid beetles (Noda and Koizumi *et al*., 2003). What properties could we use to identify a novel sterol-C7-reductase? It might be expressed in the prothoracic gland and adult reproductive tissues, consistent with the pattern of Neverland expression in most insects, though as it acts immediately following steroid dealkylation – and possibly even before DHCR24, given evidence of 7-dehydrodesmosterol synthesis in *Tribolium confusum* (Svoboda *et al*., 1975) – expression in the gut is also a possibility. Considering expression patterns therefore does less than might be hoped to narrow down the list of candidates, justifying a consideration of the enzyme families in which sterol-C7-reductase activity is most likely to evolve. Of course, another Rieske-domain enzyme could have assumed Neverland’s role; this possibility is more difficult to test than it might appear, due to poorly conserved regions within the Rieske domain (Lebrun *et al*., 2006). A small number of cysteine and histidine residues crucial to binding the iron-sulfur cluster are key to recognising Rieske-domain proteins. Querying the *T. castaneum* protein set with the Pfam Rieske-domain oxygenase profile returned only sequences annotated as ‘AIF3’, ‘cytochrome b-c1’ and ‘ubiquinol-cytochrome C’ reductase as hits. These are conserved roles for Rieske domain oxygenases (Susin *et al*., 1999; Iwata *et al*., 1998) – while ‘cytochrome b-c1’ and ‘ubiquinol-cytochrome C’ are alternate names for the same enzyme, orthologs of both *T. castaneum* hits can be found in non-coleopteran insects (e.g. *Bombyx mori* LOC101736488 and LOC134198821). It is therefore unlikely that any of these enzymes has acquired a novel role as a Neverland replacement.

Several other candidate enzyme families deserve consideration on the basis of observed sterol desaturase activity among their members. The fatty acid hydroxylase superfamily is a relatively simple analytic target, as the number of such enzymes in arthropods is small; three are found in *D. melanogaster*, namely CG1998, CG11162 and fa2h. The last of these, as its name suggests, is a fatty acid 2-hydroxylase (Carvalho *et al*, 2010), though the other two have not been functionally characterised. Orthology analysis suggests CG1998 and CG11162 also have fatty acid substrates (Shen *et al*., 2010), and expression of CG1998 in the head (Aradska *et al*., 2015) is consistent with this hypothesis given synthesis of fatty acid amides in the *D. melanogaster* head (Jeffries *et al*., 2014). Expression of CG11162 is restricted to the midgut, naturally an organ essential for the uptake of fatty acids derived from dietary lipids. Only two genes containing fatty acid hydroxylase domains are present in *T. castaneum*, one akin to fa2h and the other to CG1998/CG11162; there is thus no obvious sign of a novel sterol oxygenase among the fatty acid hydroxylase superfamily in beetles. Until the discovery of Neverland, it was generally accepted that sterol-C7-desaturation was catalysed by a cytochrome P450 (e.g. Grieneisen *et al*., 1993), and it is certainly possible that this is in fact the case in beetles. However, given the large number of cytochrome P450s present in many beetles (e.g. Zhu *et al*., 2013), it would be useful to cut down the number of candidates under consideration. Cytochrome P450 desaturases acting on the sterol side-chain are known from fungi (CYP61, Skaggs *et al*., 1996) and plants (CYP710, Morikawa *et al*., 1996), but neither of these cytochrome P450 lineages is represented in arthropods. Various sterol hydroxylases of course exist in arthropods – they catalyse many of the reactions of ecdysteroid synthesis. However, there are no obvious novel beetle genes in the CYP2 and mitochondrial cytochrome P450 clades to which established ecdysteroidogenic cytochrome P450s belong; while CYP353 was originally (Zhu *et al*., 2013) described from *Tribolium castaneum*, it is not confined to beetles. There are thus no clearly outstanding candidate sterol-C7-desaturases among beetles at present, with further consideration of transcriptome evidence clearly required.

Finally, it could be asked how important the loss of Neverland is – that is, in groups where a novel sterol-C7-reductase has arisen, should we expect to see identifiable effects of this substitution on sterol metabolism? Evidence from beetles suggests that this may in fact be the case; the balance between cholesterol and 7-dehydrocholesterol in beetles is often tilted more towards the latter than is typical for insects (e.g. Svoboda and Lusby, 1994). This finding may suggest a more efficient sterol oxygenase than Neverland, though observation of sterol kinetics in a broader range of beetles would be needed for confirmation, and of course it is unclear whether similar effects should be expected in any other groups which have developed alternative sterol-C7-reductases.

#### 3.2.2. Neverland Blooms

The hypothesis that loss of Neverland is typically driven by ergosterol-rich diets is in tension with the presence of two or three Neverland paralogs in many fungus-cultivating ants (Attini), with the occurrence of a single Neverland enzyme in some species reflecting secondary loss. How might this odd finding be explained? An examination of the sterols ingested by *Acromyrmex octospinosus* determined not only that Δ5,Δ7-sterols dominate dietary sterol intake in this species, but that sterols lacking both double bonds were selectively excreted, consistent with a lack of sterol desaturase activity (Maurer *et al*., 1992). One seemingly promising explanation for these findings would be a functional shift in ancestral attine Neverland to a role in which copy number increase is selectively favoured, such as detoxification. However, while one clade of attine Neverland paralogs is clearly evolving more rapidly than the other, even the faster of the two appears to be evolving only modestly more quickly than Neverland orthologs in other ants (Figure 4). The example of *D. pachea* Neverland suggests that catalytic activity in this enzyme can be altered with only a small number of amino acid changes, but Contrast-FEL analysis finds no individual sites under differential selection between the combined attine Neverland and non-attine ant Neverland groups after correction for false discovery rate. The same is true if the more rapidly-evolving attine clade is compared to all other myrmicine Neverland orthologs. Functional characterisation of these paralogs will be needed to resolve the paradox of Neverland expansion in an ergosterol-dependent clade.

**Figure 4.**
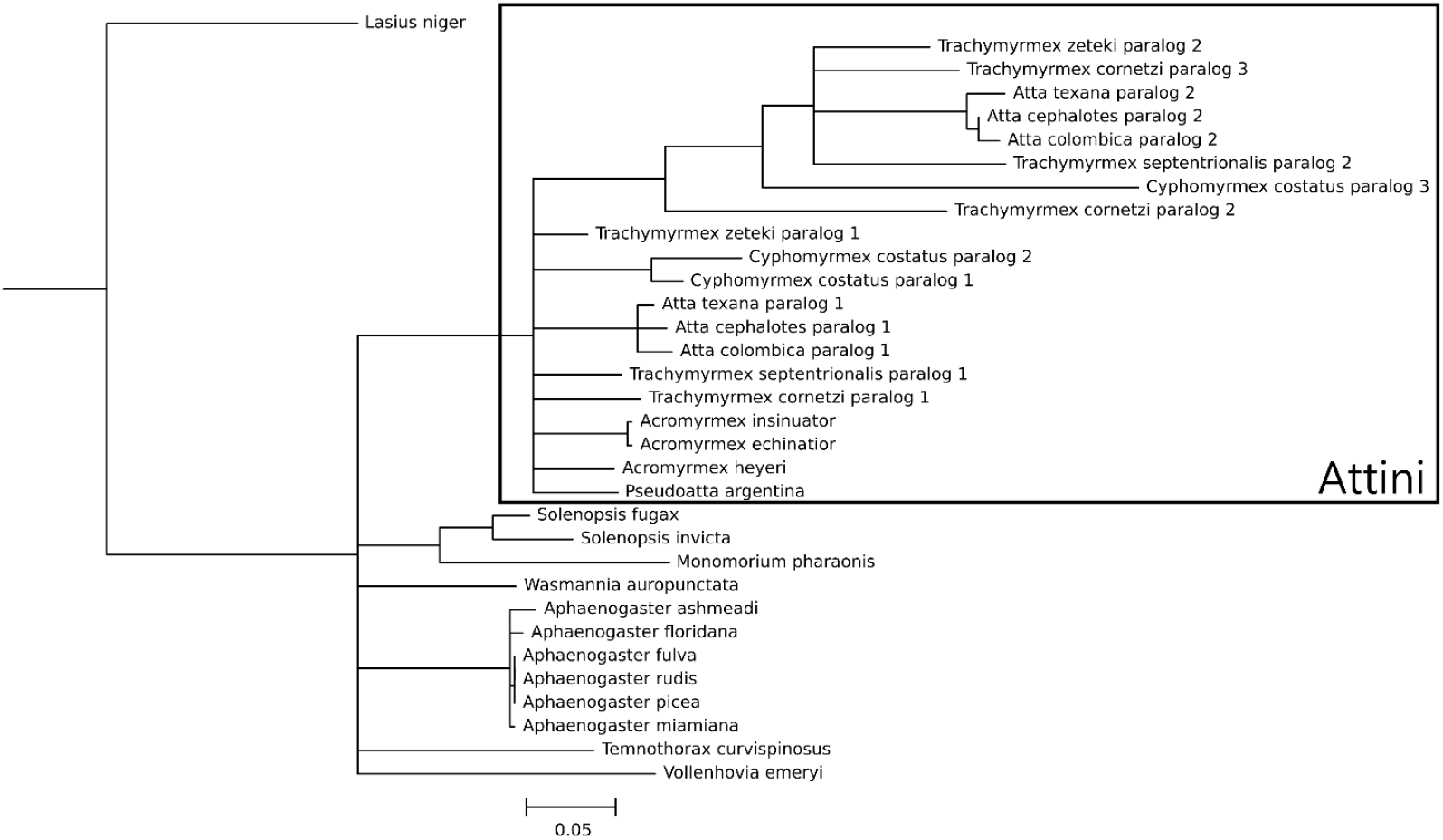
Maximum likelihood tree of Neverland orthologs in Myrmicinae. All myrmicine species in our dataset (Supplementary Table 1) from which full-length amino acid sequences could be determined are included. *Lasius niger* (Formicinae) was set as an outgroup following the phylogeny of Ward *et al*. (2015).

While not bearing so directly on questions concerning the role of ergosterol as a driver of Neverland evolution, other Neverland duplications present additional mysteries. In both chelicerates (especially mites) and springtails, eight or nine copies of Neverland are not unusual, and more can sometimes be seen – sixteen genes encoding Neverland orthologs without obvious inactivating mutations are present in the genome assembly of the orbatid mite *Achipteria coleoptrata*. It seems implausible in the extreme that all these enzymes are required for the conversion of cholesterol to 7-dehydrocholesterol in the context of ecdysteroid synthesis; as the rate-limiting step of ecdysteroid synthesis is thought to occur within the Black Box (e.g. Niwa *et al*., 2010), it is not clear that a duplication of Neverland would increase the rate of ecdysteroid synthesis. Even if it were to do so, however, an eighth or ninth copy, in the absence of duplication of any other ecdysteroid synthesis enzyme, could surely have no effect on the dynamics of the pathway.

What other role, though, might these paralogs play? Specialisation on different dietary sterols might be invoked, but the affected species do not appear to have exceptionally diverse diets, and in any case most sterol structural variation involves the side-chain, and Neverland’s site of activity is quite distant from the side-chain. Another possibility is that many of the Neverland paralogs act on non-sterol substrates. For example, as aromatic compounds are common substrates of Rieske-domain oxygenases, the synthesis of benzenoid defensive compounds by the mite *Archegozetes longisetosis* (Brückner *et al*., 2020) might involve the activity of some of its eight Neverland paralogs; some unusual defensive compounds synthesised by collembolans are also known (Möllerke and Schultz, 2024). These are of course speculative suggestions, but could be tested by examination of paralog expression patterns and ultimately functional characterisation of the Neverland paralogs.

The evolution of both DHCR24 and Neverland in Arthropoda reflects the complexities of their origin in species capable of *de novo* sterol synthesis. Beyond this point in ecdysteroid synthesis, we reach the Black Box – a series of reactions apparently unique to arthropods and poorly understood due to the instability of its intermediates. However, while knowledge of vertebrate sterol metabolism cannot assist us in understanding the Black Box, a comparison with a closer relative has the potential to do so.

### 3.3. Unshrouding the Black Box

The parallel between 3-hydroxyl-sterol-reduction in arthropod and nematode sterol hormone synthesis led us to examine the relationships between *C. elegans* and *D. melanogaster* short-chain dehydrogenase/reductases. While these enzymes do not show generally one-to-one orthology, DHS-16 and Shroud belong to a clade containing two other enzymes from each phylum, namely *C. elegans* DHS-2 and DHS-20 and *D. melanogaster* CG8888 and CG13377 (Figure 5). While this relationship between DHS-16 and Shroud provides some support for Shroud as a 3-hydroxy-sterol-reductase, the question of the function of other members of this group ought to be considered.

**Figure 5.**
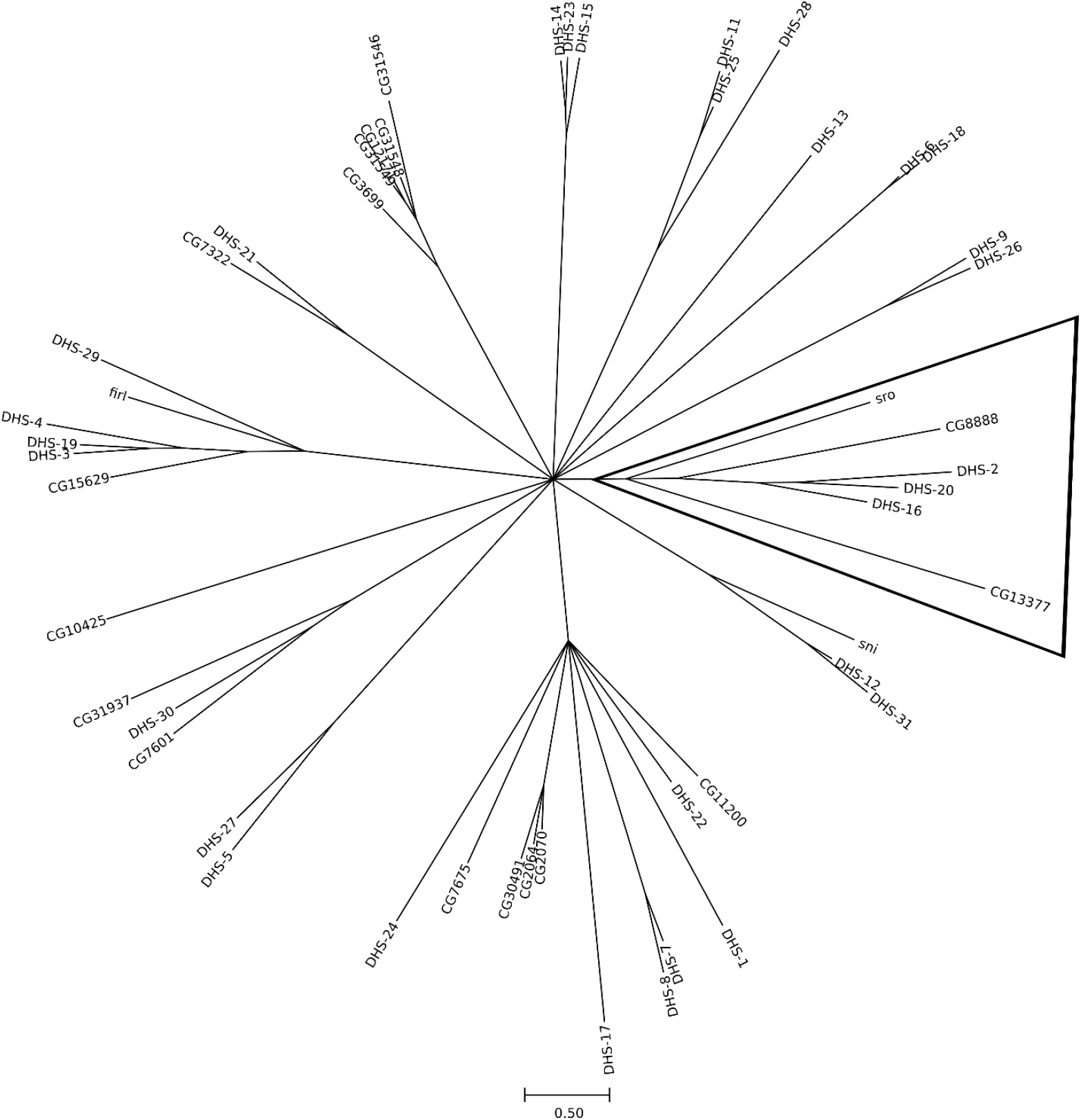
Phylogeny of short-chain reductases in *D. melanogaster* and *C. elegans*. The clade containing Shroud and DHS-16 is highlighted.

CG8888 is expressed in the corpora allata, best known for their role in the synthesis of juvenile hormone bisepoxide (the presence of which determines whether a moult is followed by another larval stage or the onset of pupariation). A short-chain reductase has been identified as catalysing oxidation of farnesol in juvenile hormone synthesis in *Aedes aegypti* (Mayoral *et al*., 2009), but CG8888 is not among the closest relatives of this enzyme in *D. melanogaster*. This fact does not necessarily exclude the possibility that CG8888 encodes a farnesol oxidase, as none of the six short-chain reductases which cluster with *A. aegypti* farnesol oxidase shows specific expression in the corpora allata. The enzyme encoded by CG8888 might also act in the subsequent step of juvenile hormone synthesis, the conversion of farnesal to farnesoic acid. Again, this step of the pathway is better understood in mosquitoes, with a fatty aldehyde dehydrogenase found to catalyse this reaction in *A. aegypti* (Rivera-Perez *et al*., 2013). This gene has a single clear ortholog in *D. melanogaster, Aldh-III*; it is broadly expressed, and thus its product clearly does not act solely in juvenile hormone synthesis. A more targeted analysis of *Aldh-III* will be needed to determine whether it is responsible for the synthesis of farnesoic acid. Of course, it is not necessarily safe to assume that every metabolic enzyme specifically expressed in the corpora allata must act directly in the synthesis of juvenile hormone, but the candidacy of CG8888 for a role in the early stages of this process nonetheless deserves further investigation.

CG13377 is not expressed in any endocrine tissue, instead being active in the embryonic central nervous system. Functional information about the CG13377 product is limited, but P-element insertion upstream of CG13377 causing transcription upregulation has been found to drive increased aggression in adult males (Edwards *et al*., 2009). Increased adult male aggression is also associated with lowered CYP6A20 levels (Dierick and Greenspan, 2007); the substrate of CYP6A20 has not been identified, but deficient degradation of male pheromones has been proposed as mediating this effect on aggression (Dierick and Greenspan, 2007). The pheromone 11-*cis*-vaccenyl acetate has been described as promoting aggression in male *D. melanogaster* (Wang and Anderson, 2010), and its receptors are co-expressed with CYP6A20 in the trichoid sensilla (van der Goes van Naters and Carlson, 2007). Linking the aggressive phenotype associated with CG13377 to this pheromone would require the putative reductase activity of this enzyme to either increase the potency of 11-*cis*-vaccenyl acetate or to block the action of enzymes degrading the pheromone, likely including CYP6A20.

While the functions of both CG8888 and CG13377 remain unclear, there is no evidence that either acts on a sterol, providing some evidence against sterol metabolism as an ancestral role for this clade of short-chain reductases. Another factor which should be considered in evaluating a role for Shroud as a steroid-3-hydroxyreductase is the fact that 3-dehydroecdysteroids have low levels of hormonal activity, and reversal of 3-hydroxy-reduction is therefore a standard part of ecdysteroid synthesis (though its timing relative to other reactions is somewhat variable). It would seem reasonable to expect that an ultimately reversed reaction would at least occasionally be lost among arthropods. Shroud in fact seems to be quite stable; its best-supported loss is found in barnacles, which have also lost several other essential ecdysteroid synthesis genes (Dermauw *et al*., 2020).

What to make of this loss is unclear – barnacles have retained other genes acting in ecdysteroid synthesis, and while few direct examinations of barnacle ecdysteroids have been conducted, and they are a challenging group in which to control for dietary intake, several studies have reported the presence of 20-hydroxyecdysone in barnacles (Bebbington *et al*., 1997; Morgan *et al*., 2005). While the parallel to nematode dafachronic acid synthesis remains intriguing, evidence for Shroud’s specific role in ecdysteroid synthesis remains equivocal at best.

Duplications of Shroud appear sporadically, but are rarely shared across multiple currently sequenced species. A clear exception is provided by crickets; of the five cricket genome assemblies publicly available at the time of writing, only one showed no sign of Shroud duplication. Analysis of the ecdysteroids produced by *Acheta domesticus* (which shows evidence of having six copies of Shroud, though the fragmentation of its genome means pseudogenisation cannot be ruled out) does not suggest any oddities in ecdysteroid synthesis – ecdysone and 20-hydroxyecdysone are predominant (Dinan, 1997). A reasonable explanation for these findings would be development of functions for some of the cricket Shroud orthologs outside ecdysteroid synthesis, though this is difficult to test with such a paucity of genomes.

## 4. Identification of other Black Box Enzymes

Following the example of the Shroud analysis above, could we also use homology to identify the enzymes responsible for other Black Box reactions? If we are willing to take a broad interpretation of relevant parallels, there might seem to be a wealth of information available; for instance, 6-hydroxylation of steroid hormones in the vertebrate liver by CYP3A4 (Waxman *et al*., 1988) could be interpreted as analogous to the first step of steroid-6-oxidation within the Black Box. The question, of course, is where such an identification is meaningful. CYP3A4 is a relatively promiscuous detoxification enzyme, and there are substantial structural differences between vertebrate steroid hormones and Black Box intermediates (most obviously, the former lack a side chain). Perhaps most importantly, the almost entirely independent developments of the cytochrome P450 superfamily in arthropods and vertebrates (e.g. Markov *et al*., 2009) means that steroid-6-oxidation must have evolved independently in each group.

An alternative approach would be to consider in more detail enzymes already associated with the Black Box. CYP307 is of particular interest due to its apparent propensity towards duplication (e.g. Ono *et al*., 2006); this may reflect activity in the rate-limiting step of ecdysteroid synthesis (which in turn might mean that these duplications are particularly relevant in allowing use of varied substrates). Another possibility is that CYP307 ancestrally catalysed two distinct Black Box reactions (e.g. steroid-6- and 14-hydroxylation), with CYP307 duplications resulting in subfunctionalisation, though the fact that CYP307 paralogs tend to have non-overlapping expression patterns (e.g. Sztal *et al*., 2007; Hentze *et al*., 2013) argues against this interpretation. However, even if CYP307 enzymes only catalyse one of the Black Box reactions, there is still no shortage of enzymes to which roles in the Black Box have been attributed. Two other cytochrome P450s, CYP6T3 and CYP6U1, have been associated with the *D. melanogaster* Black Box on the grounds that interrupting synthesis of these enzymes in the prothoracic gland perturbs ecdysteroid synthesis (Ou *et al*., 2011). However, both these enzymes would make atypical ecdysteroidogenic cytochrome P450; they are not conserved across insects (Dermauw *et al*., 2020), and where other ecdysteroidogenic cytochrome P450s belong either to the mitochondrial or CYP2 clans, CYP6T3 and CYP6U1 are members of the CYP3 clan. Mammalian CYP3A4, mentioned above, also belongs to the CYP3 clan, and it is thus certainly possible for such enzymes to act on sterol substrates. However, such activity seems to occur in the context of detoxification in mammals, and many close relatives of CYP6T3 and CYP6U1 have well-established detoxification activity in *D. melanogaster* (e.g. Daborn *et al*., 2001). The expression patterns of *Cyp6t3* and *Cyp6u1* are also atypical for ecdysteroid synthesis genes; Chung *et al*. (2009) indicate that the larval gonads are the major expression site of *Cyp6u1*, while *Cyp6t3* was found to be expressed only in adults. Finally, Shimell and O’Connor (2022) have reported that CRISPR-generated *Cyp6t3* null alleles are homozygous viable and show normal developmental timing, which suggests that loss of *Cyp6t3* does not grossly perturb ecdysteroid synthesis. It seems likely that Ou *et al*.’s (2011) results are attributable to off-target effects of RNA interference; generation of *Cyp6u1* null mutants would clarify whether such effects can also explain Christesen *et al*.’s (2017) results.

On the whole, then, the Black Box remains poorly understood. In particular, we lack plausible candidates for C6 oxidation or C4(5) reduction, both clearly highly conserved reactions. Given the example of other ecdysteroid synthesis enzymes, the most plausible assumption is that those responsible for catalysing these reactions are well conserved - but if that is the case, why have they proven so elusive in comparison to the ecdysteroidogenic cytochrome P450s (Warren *et al*., 2002; Petryk *et al*., 2003; Warren *et al*., 2004; Ono *et al*., 2006)? Increasingly diverse genomic data and the development of new analytic tools such as computational protein structural modelling (Jumper *et al*., 2021; Varadi *et al*., 2022, Lin *et al*., 2023) may assist in the identification of further Black Box candidates, but we are still a long way from full characterisation of the ecdysteroid synthesis pathway.

## 5. Conclusion

The loss of cholesterol synthesis in the arthropod lineage might have been expected to set their hormonal development on a very different trajectory to that of vertebrates, yet in fact steroid hormones play an essential role in arthropod development. The early stages of ecdysteroid synthesis are of particular value in understanding the ancient developments in sterol metabolism which gave rise to the ecdysteroid synthesis pathway, though they are of course also shaped by more recent events. In particular, the comparative analysis reported here reveals the extent to which diet continues to shape the evolution of these early ecdysteroidogenic genes; diet was likely also crucial to the loss of cholesterol synthesis and development of steroid hormones in the arthropod lineage. While there are still many leads to pursue within Arthropoda, the study of dietary evolution and sterol metabolism in arthropod relatives which have maintained *de novo* cholesterol synthesis (namely velvet worms and tardigrades) would be an obvious boon to our understanding of the evolution of sterol metabolism – there are always more comparisons to be made.

