## Supplementary figures and images for "Before the Black Box"

### Supplementary Figure 1

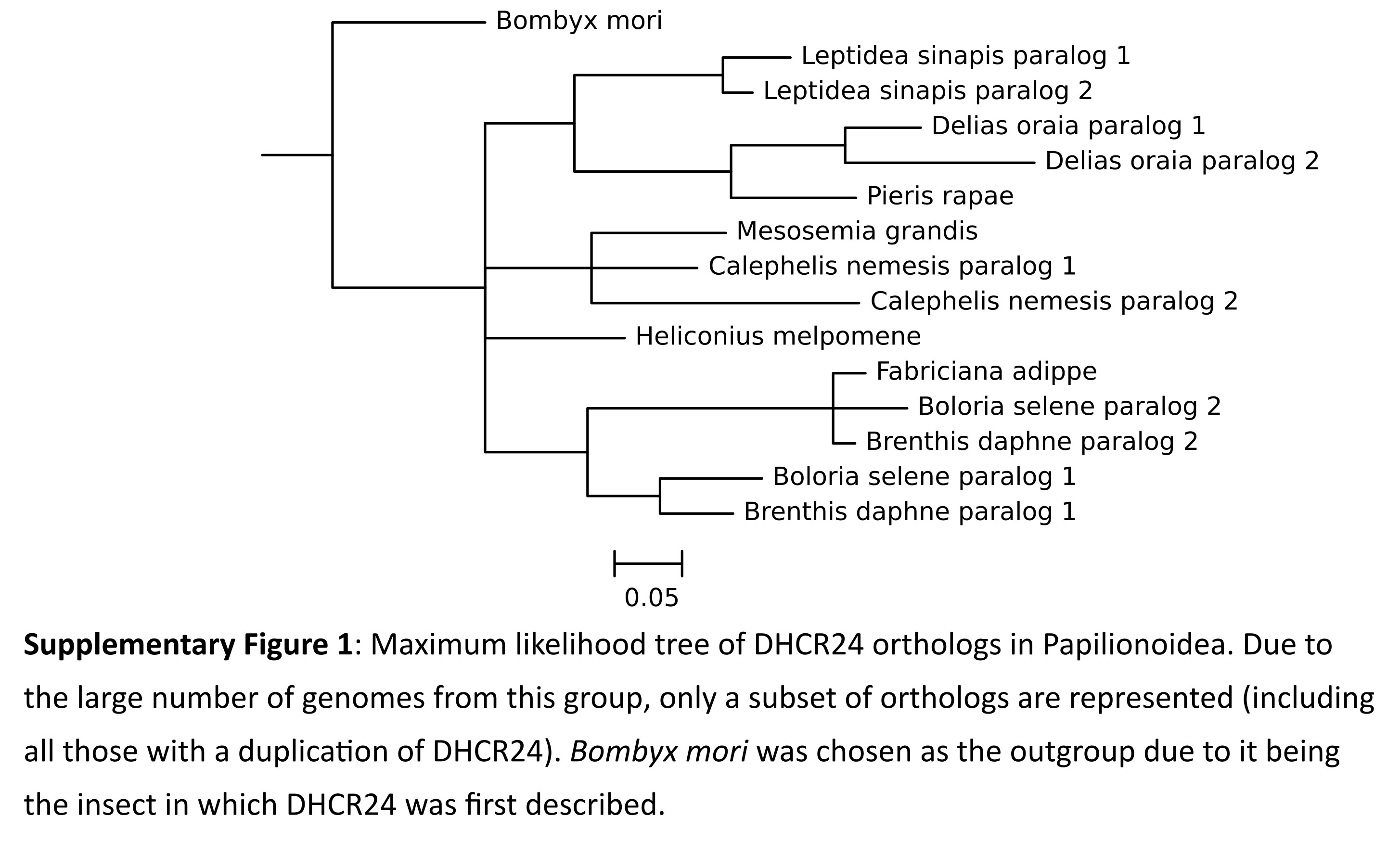

### Supplementary Figure 2

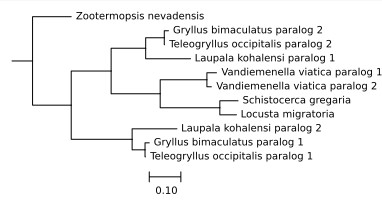
